# Active components of commonly prescribed medicines affect influenza A virus-host cell interaction: a pilot study

**DOI:** 10.1101/2021.07.11.451833

**Authors:** Aleksandr Ianevski, Rouan Yao, Eva Zusinaite, Hilde Lysvand, Valentyn Oksenych, Tanel Tenson, Magnar Bjørås, Denis Kainov

**Author notes:** Contributed equally (Aleksandr Ianevski, Rouan Yao).

## Abstract

**Background:** Every year, millions of people are hospitalized, and thousands die from influenza A virus (FLUAV) infection. Most cases of hospitalizations and death occur among elderly. Many of these elderly patients are reliant on medical treatment of underlying chronic diseases, such as arthritis, diabetes, and hypertension. We hypothesized that the commonly prescribed medicines for treatment of underlying chronic diseases can affect host responses to FLUAV infection, and thus contribute to morbidity and mortality associated with influenza. Therefore, the aim of this study was to examine whether commonly prescribed medicines could affect host responses to virus infection *in vitro*.

**Methods:** We first identified 45 active compounds of commonly prescribed medicines. Then we constructed a drug-target interaction network and identified potential implication of these interactions for FLUAV-host cell interplay. Finally, we tested the effect of 45 drugs on viability, transcription and metabolism of mock- and FLUAV-infected human retinal pigment epithelial (RPE) cells.

**Results:** *In silico* drug-target interaction analysis revealed that many drugs, such as atorvastatin, candesartan, and hydroxocobalam, could target and modulate FLUAV-host cell interaction. *In vitro* experiments showed that these and other compounds at non-cytotoxic concentrations differently affected transcription and metabolism of mock- and FLUAV-infected cells.

**Conclusion:** Many commonly prescribed drugs modulate FLUAV-host cell interactions *in silico* and *in vitro* and, therefore, could affect their interplay *in vivo*, thus, contributing to morbidity and mortality of patients with influenza virus infections.

## 1. Introduction

Influenza A (FLUAV) is the most common cause of seasonal epidemics. It is responsible for 3 to 5 million cases of hospitalizations and 250.000–500.000 deaths annually, negatively impacting public health and global economy. Although everyone is susceptible to influenza infection, most cases of hospitalizations and deaths occur among high-risk groups, which include pregnant women, young children, the elderly, and patients with immunosuppressive conditions or non-communicable diseases (NCDs) [1]. Deaths and hospitalizations from influenza infection are often linked to underlying conditions such as asthma, diabetes, cardiovascular, and chronic kidney diseases. For instance, influenza infection may elevate inflammation and contraction of inflamed and swollen airways, leading to severe asthma attacks and worsening of the patients’ asthma symptom control [2].

FLUAV belongs to the family *Orthomyxoviridae*. Its genome comprises of 8 single-stranded viral RNA segments (vRNA) of negative polarity. Each segment interacts with three viral polymerase subunits (PA, PB1 and PB2) and nucleoproteins (NP) to make eight viral ribonucleoproteins (vRNPs). vRNPs are surrounded by a capsid. The capsid is composed of matrix protein 1 (M1) and a lipid membrane derived from the host cell. The membrane is embedded with hemagglutinin (HA), neuraminidase (NA), and matrix protein 2 (M2). Other viral proteins, such as non-structural protein 1 (NS1), nuclear export protein (NEP), polymerase basic 1 F2 (PB1-F2), polymerase acidic X (PA-X), and the N-deleted version of PB1(N40) are not present in the virion and only expressed in infected cells [3].

The FLUAV replication cycle consists of entry into the host cell through endocytosis, uncoating of vRNPs, import of the vRNPs into the nucleus, transcription and replication of the viral genome, translation of viral proteins, assembly of vRNPs in the nucleus, export of the vRNPs from the nucleus, assembly virions at the host cell plasma membrane and budding [3]. When FLUAV enters the cell, pathogen recognition receptors (PRRs) recognize viral RNA (vRNA) and initiate the transcription of interferon (IFN) genes. Once transcribed, IFNs trigger the expression of IFN-stimulated genes (ISGs) in an infected cell and, when secreted, in neighboring non-infected cells. ISGs encode different antiviral proteins including interleukins (ILs), C-X-C and C-C motif chemokines (CXCLs and CCLs), and other cytokines which recruit immune cells to the site of infection. ISGs also encode RNases, which degrade vRNAs in infected cells. NS1 hinders the cellular IFN response by binding with vRNA, cellular DNA and other factors [4-6]. If large amount of vRNA or its replication intermediates is accumulated in the cells, apoptosis is initiated. For this to occur, PRRs recognize vRNA and transduce signals to B-cell lymphoma xL (Bcl-xL) protein [7, 8]. Bcl-xL releases pro-apoptotic proteins to initiate mitochondrial outer membrane permeabilization (MoMP), which results in infected cell death. If IFN or apoptotic pathways are inhibited or altered, virus replication can be accelerated, and infection worsened [3].

FLUAV also relies on its ability to exploit multiple cellular factors and pathways to complete replication [9, 10]. For example, cellular vATPase acidifies the interior of late endosomes. This activates cellular serine proteases that cleave HA and mediate fusion of viral and endosomal membranes. This triggers the release of vRNPs [11]. Cellular cyclin-dependent kinases are required for vRNA transcription and free cellular nucleotides are used by viral polymerase to produce vRNA [12]. FLUAV also hijacks PI3K/mTor/Akt-mediated autophagy to produce free amino acids for synthesis of viral proteins [13]. Virus assembly and budding also depends on elements of host lipid metabolism, including de novo synthesis of cholesterol [14].

These cellular factors could be targeted or altered to inhibit virus replication. For example, the anticancer drug saliphenylhalamide (SaliPhe) targets vATPase, thus inhibiting FLUAV at the entry stage [11]. Likewise, the anticancer drugs flavopiridol and gemcitabine target CDKs and nucleotide metabolism, which inhibit viral RNA transcription and replication, respectively [12, 15]. Cholesterol-lowering statins inhibit FLUAV assembly and budding [16]. Because the complex nature of influenza infection involves many aspects of host biology, we hypothesized that some of these factors could be targeted by commonly prescribed medicines, and that this could modulate virus-host interaction and affect morbidity and mortality associated with FLUAV infection. Here, we present results from our pilot study of 45 active components of the most prescribed medicines. We showed how these agents affect FLUAV-host cell interaction *in silico* and *in vitro*.

## 2. Materials and Methods

### 2.1. Compounds, cells, and viruses

To identify the most dispensed medicines in Central Norway in 2019, we searched the Norwegian Prescription Database (www.norpd.no), which contains drugs, Anatomical Therapeutic Chemical (ATC) codes, and daily defined dosage (DDD). As search criteria, we included all age groups and both sexes. Table S1 lists 45 active compounds of the most dispensed medicines, their suppliers and catalogue numbers. To obtain 10□mM stock solutions, compounds were dissolved in dimethyl sulfoxide (DMSO, Sigma-Aldrich, Steinheim, Germany) or milli-Q water, depending on solubility. The solutions were stored at −80 °C until use. Human telomerase reverse transcriptase-immortalized retinal pigment epithelial (RPE, ATCC MBA-141) cells were grown in Dulbecco’s Modified Eagle’s F12 medium (DMEM-F12; Gibco, Paisley, Scotland) supplemented with 100 U/mL penicillin and 100 ug/ml streptomycin mixture (Pen/Strep; Lonza, Cologne, Germany), 2 mM L-glutamine, 10% heat-inactivated fetal bovine serum (FBS; Lonza, Cologne, Germany), and 0.25% sodium bicarbonate (Sigma-Aldrich, St. Louis, USA). Human influenza A/WSN/33(H1N1) virus was generated using the eight-plasmid reverse genetics system as described previously [17].

### 2.2. Cell viability assay

Approximately 4 × 10^4^ RPE cells were seeded per well in 96-well plates. The cells were grown for 24 h in DMEM-F12 medium containing 10% FBS, and Pen/Strep. The medium was replaced with DMEM-F12 medium supplemented with 0.2% bovine serum albumin (BSA), 1 μg/mL TPSK-trypsin and 2 mM L-glutamine. The compounds were added to the cells at seven different concentrations in 3-fold dilutions starting from 100 μM. Saliphenylhalamide (SaliPhe) and ABT-263 were used as controls [7, 8, 11, 15]. RPE cells were infected with FLUAV at multiplicity of infections (moi) of 1 or mock. After 48 h of infection, the medium was removed from the cells. The viability of virus- and mock-infected cells were measured using Cell Titer Glow assay (CTG; Promega, Madison, USA). The luminescence was read with a PHERAstar FS plate reader (BMG Labtech, Ortenberg, Germany).

The half-maximal cytotoxic concentration (CC_50_) and the half-maximal effective concentrations (EC_50_) for each compound were calculated using GraphPad Prism software version 7.0a. CC_50_ were calculated based on curves obtained on mock-infected cells after non-linear regression analysis with a variable slope. The EC_50_ were calculated based on the analysis of viability of infected cells by fitting drug dose–response curves using four-parameter (4PL) logistic function f(x):

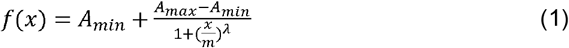

where f(x) is a response value at dose x, A_min_ and A_max_ are the upper and lower asymptotes (minimal and maximal drug effects), m is the dose that produces the half-maximal effect (EC_50_ or CC_50_), and λ is the steepness (slope) of the curve. A relative effectiveness of the drug was defined as selectivity index (SI = CC_50_/EC_50_).

### 2.3. Transcriptomics analysis

We infected RPE cells with FLUAV at moi 1. After 8 h we isolated total RNA using a RNeasy Plus minikit (Qiagen). 384 TruSeq Stranded mRNA libraries were prepared in 96 sample batches. Sequencing was done on HiSeq (HSQ-700358) instrument (set up: SR 1 x 70 bp + dual index 8 bp) using HiSeq Rapid SR Cluster Kit v2 sequencing kit, RapidRunV2 flow cell (up to 300M reads per flowcell), RTA version: 1.18.64. For the viral genome, reads were aligned using the Bowtie 2 software package version 2.3.4.1 to the reference influenza A/WSN/1933. Sequence alignments were converted to Binary alignments using SAMtools version 1.5. The number of mapped and unmapped reads that aligned to each viral gene were retrieved with SAMtools idxstats. For the human genome, reads were aligned using the Bowtie 2 software package version 2.3.4.1 to the reference human GRCh38 genome. The number of mapped and unmapped reads that aligned to each host gene were obtained with featureCounts function from Rsubread R-package version 2.10.

### 2.4. Metabolomics analysis

We infected RPE cells with FLUAV at moi 1. After 24 hours, we collected the cell culture medium. We analysed polar metabolites as described previously [18].

### 2.5. Bioinformatics analysis

Cellular targets of drug-protein interaction were visualized using the STITCH web-tool [19]. Predicted interaction sources were excluded from the network visualization, while the line thickness was set to indicate the strength of data support.

Transcriptomics and metabolomics data were log2 transformed for linear modelling. The heatmaps were generated using the pheatmap package (https://cran.r-project.org/web/packages/pheatmap/index.html) based on log2-transformed profiling data. Gene (GSEA) and metabolite (MSEA) set enrichment analysis tools were used to retrieve pathways (http://software.broadinstitute.org/gsea/index.jsp; https://www.metaboanalyst.ca/). Structural similarity between compounds was calculated using ECPF4 fingerprints and the Tanimoto coefficient.

## 3. Results

### 3.1. Structural comparison of 45 active components of commonly prescribed drugs

We selected 45 most dispensed medicines from the Norwegian Prescription Database (www.norpd.no). Table S2 lists ATC-codes, DDDs and indications retrieved from the common catalog of pharmaceutical preparations marketed in Norway (www.felleskatalogen.no). Over-the-counter medicines, such as anti-inflammatory ibuprofen, paracetamol, and medicines to treat allergies, heartburn, constipation, and diarrhea were not included in our study, because they are used to treat acute conditions.

Figure 1 depicts the structural relationship between 45 active components of the commonly prescribed drugs. Drugs that appear closer together on the dendrogram are more structurally related than those appearing more distant from each other. These structural relationships can often also reveal functional relationships. For example, salmeterol and salbutamol are almost structurally identical except for the absence of a long ether side chain in salbutamol. Both salmeterol and salbutamol are β2-adrenoceptor agonists which are used as smooth muscle relaxants against asthma. Other examples of structurally and functionally similar molecules include angiotensin II receptor antagonists candesartan and losartan, which are used to treat hypertension; corticosteroids mometasone furoate and fluticasone propionate, which are commonly prescribed against allergy; and proton pump inhibitors esomeprazole and pantoprazole, which are used to treat reflux and ulcers.

**Figure 1.**
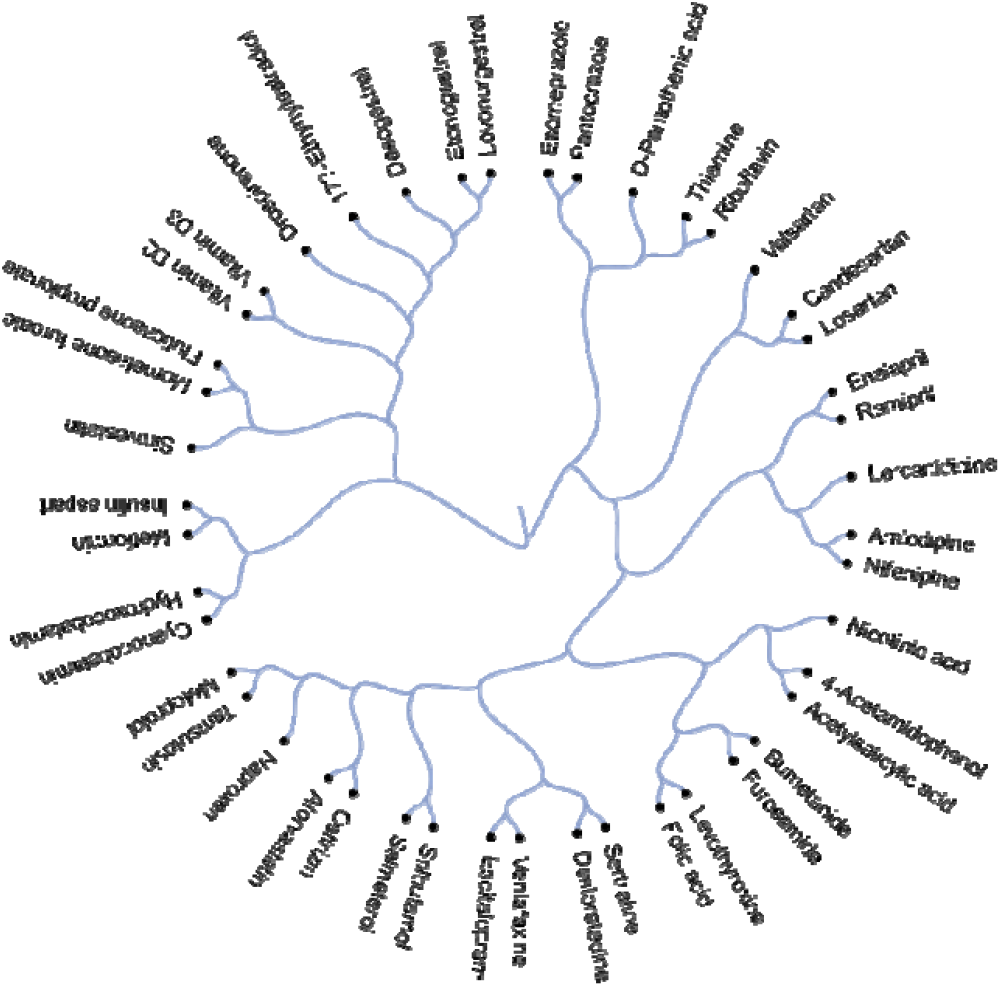
Active compounds of 45 most prescribed drugs clustered, based on their structural similarity, calculated by ECPF4 fingerprints and the Tanimoto coefficient.

### 3.2. Cellular targets of active components of the most prescribed drugs and their potential effect on FLUAV-host cell interact on

We constructed an interaction network of the 45 active components connected to direct and downstream targets (Fig. 2). The edge, width, and color darkness indicate the degree of data support for the connection. We found several targets which are associated with FLUAV replication. These include CXC chemochine receptor type 4 (CXCR4), albumin presursor (ALB), histamine receptor H1 (HRH1), Ras homolog family member A (RHOA), alpha-2B adrenergic recpetor (ADRA2B), puring nucleoside phosphorylase (PNP), and metabolism of cobalamin associated B (MMAB) proteins, which are targeted by niacin, losartan, ramipril, acetylsalicylic acid, thyroxine, valsartan, cetirizine, citalopram, atorvastatin, simvastatin, metoprolol, setraline, candesartan and hydroxocobalam [9, 10]. Thereby, commonly prescribed drugs could target and modulate FLUAV-host cell interactions, and therefore could affect morbidity and mortality of influenza-infected patients.

**Figure 2.**
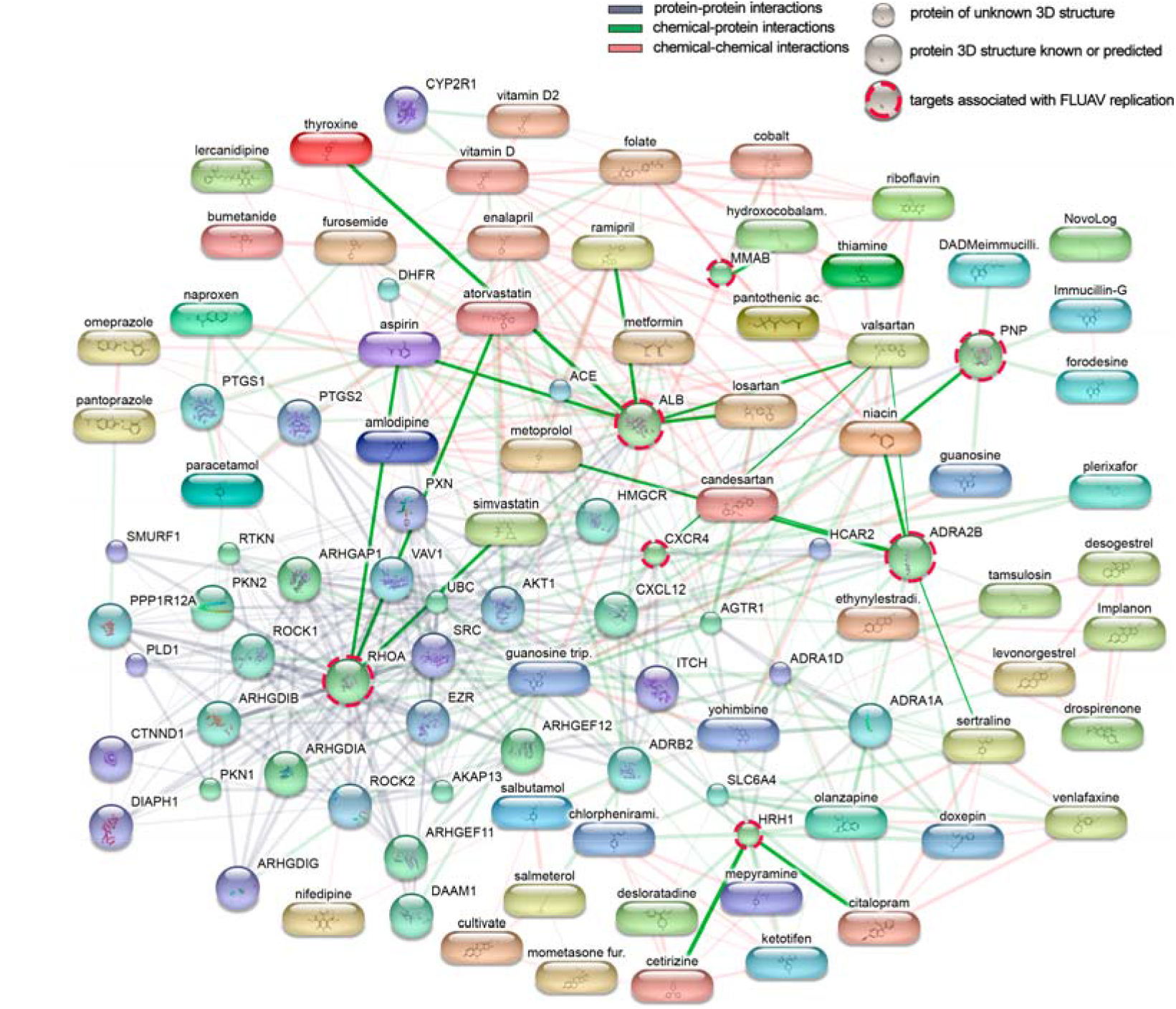
Direct and downstream cellular targets of 45 active components of commonly prescribed medicines. Targets associated with FLUAV replication are marked with red-dashed circles and interactions between them and commonly prescribed drugs are highlighted.

### 3.3. Toxicity and efficacy of active components of commonly prescribed medicines in mock- and FLUAV-infected RPE cells, respectively

In order to examine the effect of active components of commonly prescribed medicines on viability of RPE cells, the cells were treated with compounds in 3-fold dilutions at 7 different concentrations starting from 100 μM. No compounds were added to the control wells. After 48 h, cell viability was measured. The half-maximal cytotoxic concentration of each compound were determined and plotted (Fig. 3). We found that all compounds except 8 were nontoxic at the tested range of concentrations (<100 μM). The 8 counpounds that demonstrated toxicity were amlodipin (CC_50_=28,5 μM), desloratadine (CC_50_=46,7 μM), desogestrel (CC_50_=34,5 μM), salmeterol (CC_50_=29,3 μM), sertraline (CC_50_=17,5 μM), simvastatin (CC_50_=48,3 μM), vitamin D2 (CC_50_=42,1 μM), and vitamin D3 (CC_50_=48,8 μM). We also determined antiviral effecacy of the active components. For this, RPE cells were treated with compounds in 3-fold dilutions at 7 different concentrations starting from 100 μM and infected with FLUAV at a moi 1. No compounds were added to the control wells. After 48 h, cell viability was measured. Half-maximal effective concentrations of each compound were determined (Fig. 3). We found that none of the compounds rescued cells from FLUAV-mediated death.

**Figure 3.**
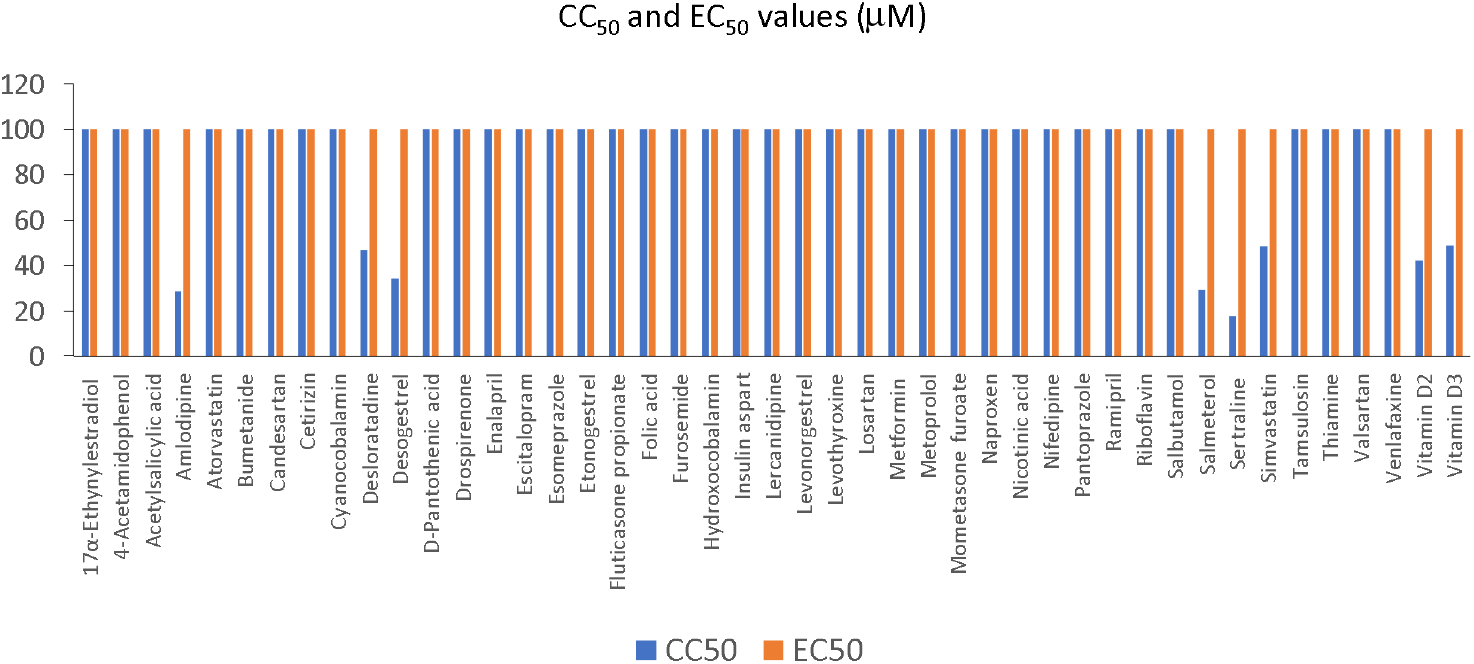
Effect of 45 active compounds of commonly prescribed drugs on the viability of mock- and FLUAV-infected RPE cells. CC_50_ and EC_50_ values are shown in blue and orange, respectively (Mean, n = 3).

### 3.4. Active components of commonly prescribed medicines affect gene expression in mock- and FLUAV-infected RPE cells

We evaluated the effects of the compounds on transcription of host genes in mock and virus-infected RPE cells. The cells were either treated with 10 μM drug or remained untreated, and either infected with mock or FLUAV (moi 1). After 8 h, we isolated total RNA and sequenced the polyadenylated fraction of RNA. We constructed a heatmap of the most variable genes of mock- and FLUAV-infected cells (Fig. 4 and 5). Almost all compounds affected transcription of host genes in both uninfected and FLUAV-infected RPE cells. Interestingly, structuraly similar drugs salmeterol and salbutamol both increased expression of phosphodiesterase 4D (PDE4D), cysteine-rich secretory protein LCCL domain-containing 2 (CRISPLD2), and prostaglandin E synthase (PTGES), and decreased expression of solute carrier family 26 member 4 (SLC26A4), family with sequence similarity 111, member B (FAM111B), and cyclin E2 (CCNE2) in non-infected cells compared to the levels found in non-treated cells, but did not affect FLUAV-activated host gene expression. Several compounds also increased expression of prostaglandin-endoperoxide synthase 2 (PTGS2), a gene known to play a role in resolution of both infectious and non-infectious inflammation, in both mock- and virus-infected cells.

**Figure 4.**
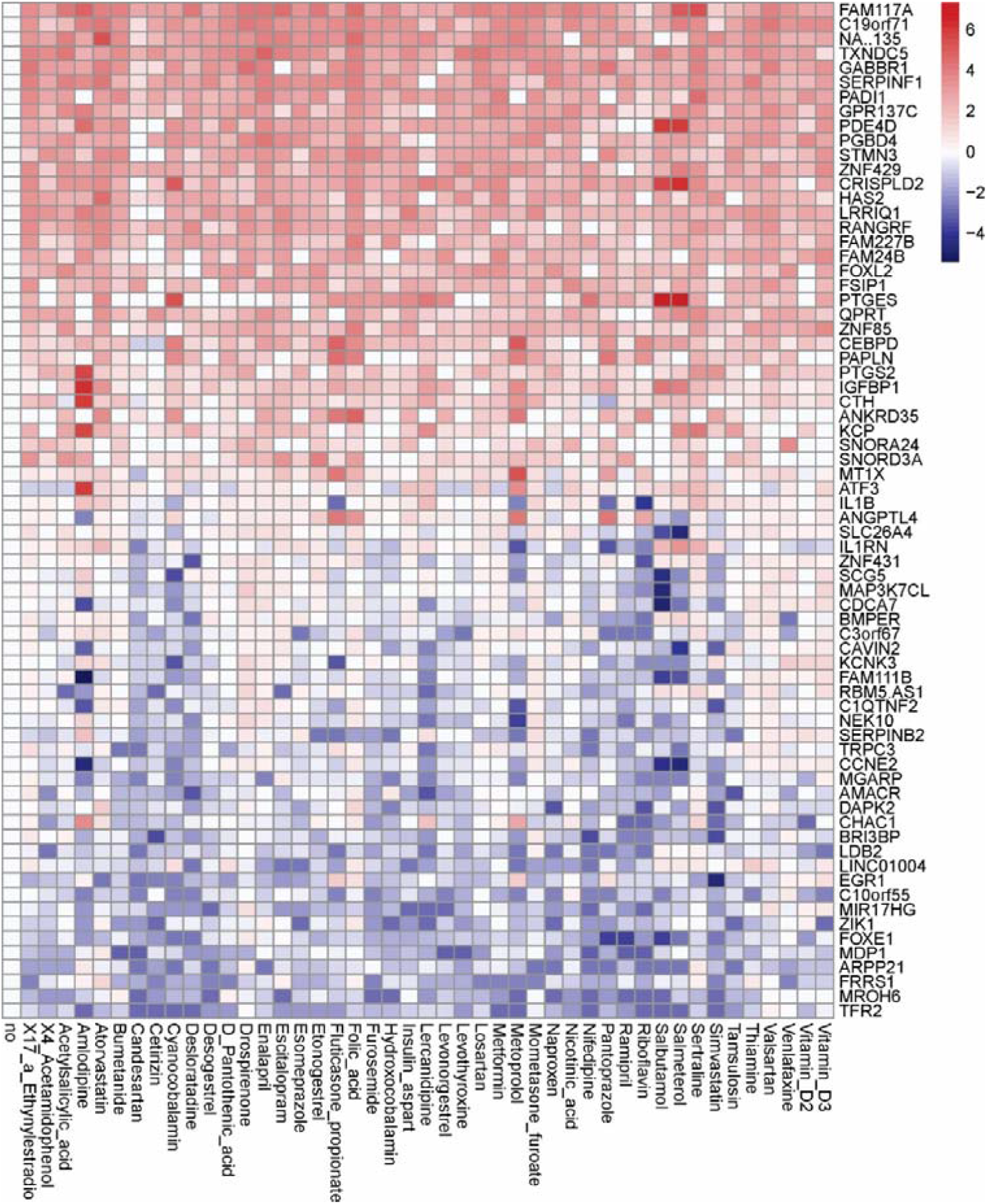
Effect of 45 compounds on polyadenylated host RNAs in RPE cells. RPE cells were treated with 10 µM compounds or remained non-treated. After 8 hours, total RNA was isolated, and a fraction of polyadenylated RNA was sequenced. A heatmap of 70 most variable mRNAs affected by treatment is shown. Rows represent gene symbols, columns represent samples. Each cell is colored according to the log2-transformed expression values of the samples, expressed as fold-change relative to the non-treated control.

**Figure 5.**
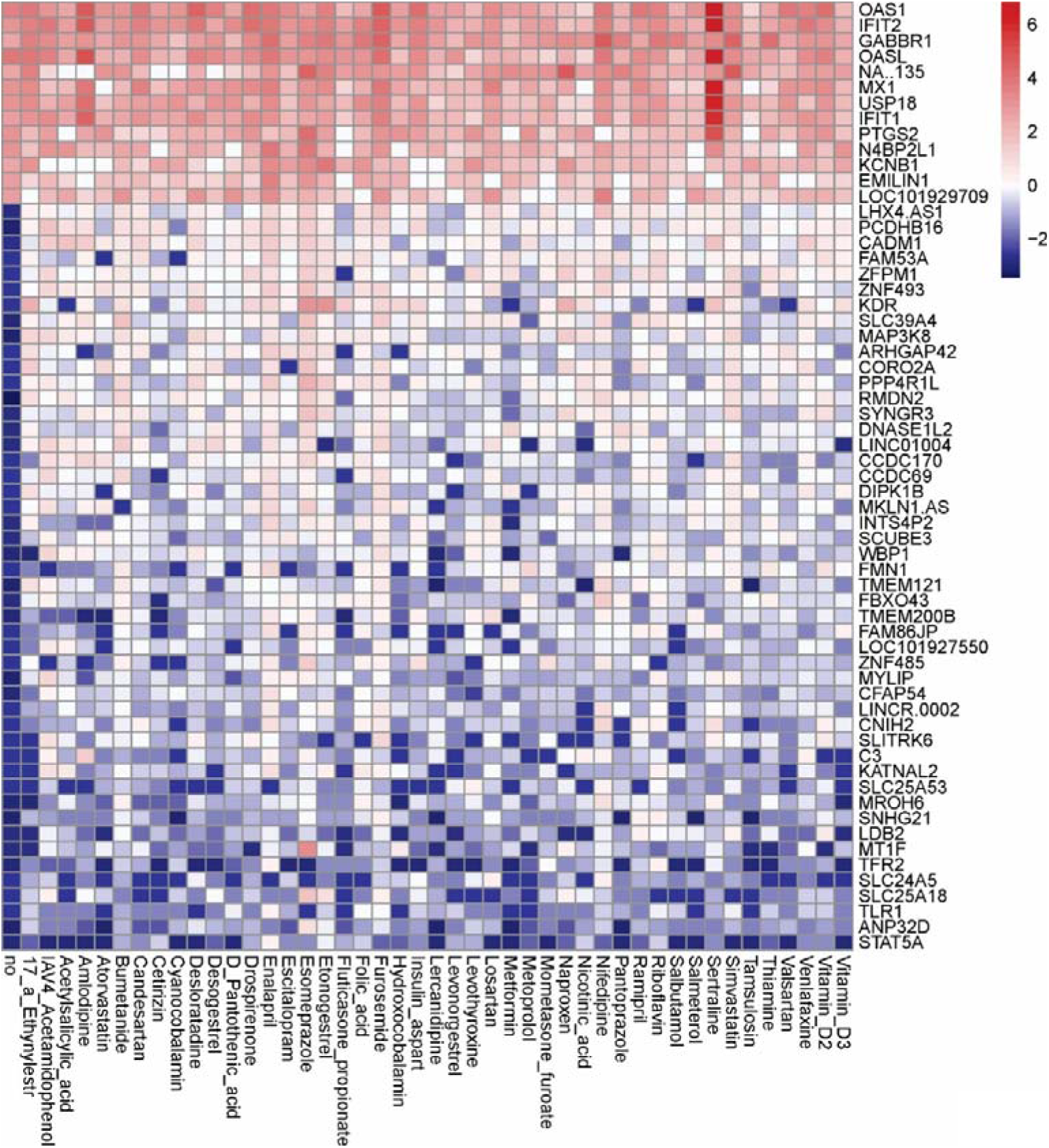
Effect of 45 compounds on polyadenylated host RNAs in FLUAV-infected RPE cells. RPE cells were treated with 10 µM compounds and infected with FLUAV at moi 1. After 8 hours, total RNA was isolated, and a fraction of polyadenylated RNA was sequenced. A heatmap of the most variable genes affected by FLUAV infection is shown (2.5 < log2FC < -2.5). Rows represent gene symbols, columns represent samples. Each cell is colored according to the log2-transformed expression values of the samples, expressed as fold-change relative to the nontreated mock-infected control.

We also analysed viral polyadenylated RNAs in drug-treated and non-treated cells (Fig. 6). Atorvastatin, cetirizine, acetylsalecytic acid, nicotinic acid, naproxen, tamsulozin, thiamine and vitamin D3 attenuated transcription of all viral RNAs (fold change >2 log2). Interesting, atorvastatin and cetirizine share structural similarlities, as does acetylsalicylic acid and nicotinic acid, indicating that there is a structure-activity relationship of drugs that decrease viral gene expression. By contrast, metformin differencially affected polyadenylated viral RNAs, increasing expression of some while decreasing expression of others. Taken together, we can see that use of these medications can indeed affect both host and viral gene expression.

**Figure 6.**
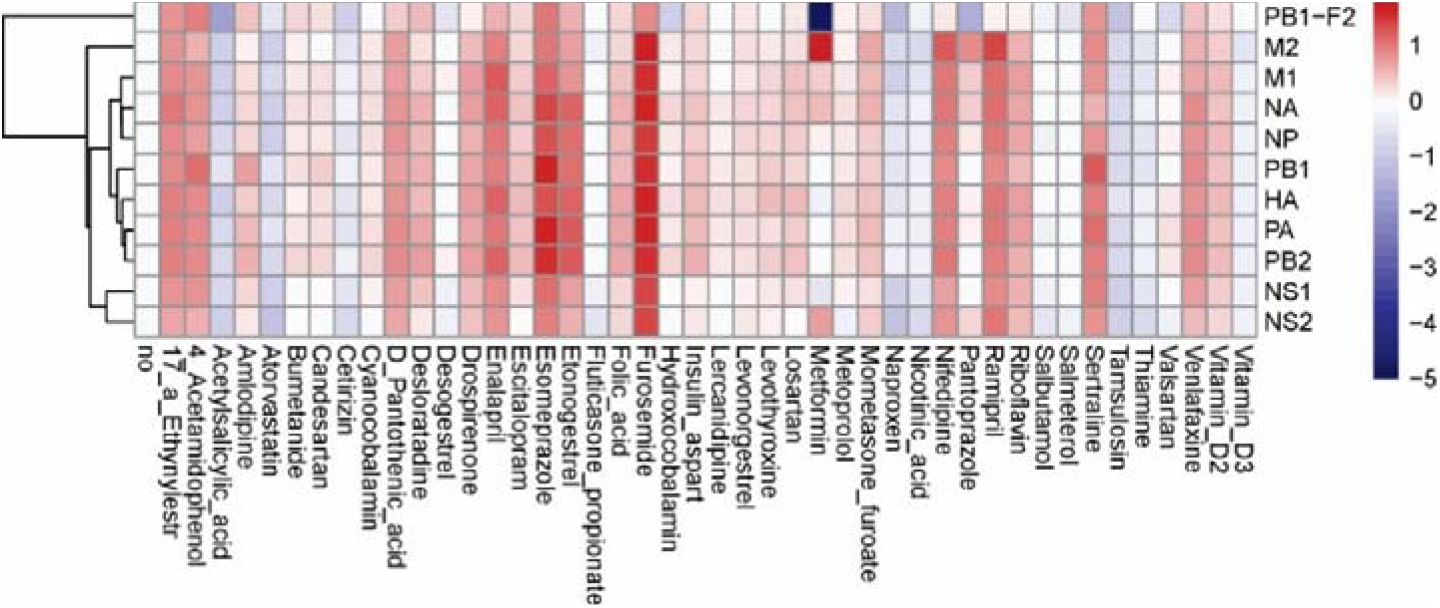
Effect of 45 compounds on polyadenylated viral RNAs in FLUAV-infected RPE cells. RPE cells were treated with 10 µM compounds or remained non-treated and infected with FLUAV at moi 1. After 8 h, total RNA was isolated, and a fraction of polyadenylated RNA was sequenced. A heatmap of viral RNAs affected by treatment is shown. Each cell is colored according to the log2-transformed expression values of the samples, expressed as fold-change relative to the nontreated FLUAV-infected control.

### 3.5. Active components of commonly prescribed medicines affect metabolism of mock- and FLUAV-infected RPE cells

Next, we evaluated the effect of the compounds on metabolism of mock and virus-infected RPE cells. The cells were either treated with 10 μM drug or remained untreated, and either mock- or FLUAV-infected (moi 1). After 24 h, the supernatants were collected and polar metabolites were analysed, and we constructed a heatmap of the most variable metabolites (Fig. 7 and 8). We found that amlodipine substantialy elevated levels of trimethylamine-N-oxide, adenine, NAD, cytosine, octanoylcarnitine, and homocysteine in the media of uninfected cells, as well as inosine and D-ribose 5-phosphate in the media of FLUAV-infected cells. By contrast, levonogestrel substantialy lowered the levels of trimethylamine-N-oxide, phosphoethanolamine, and hippuric acid in noninfected cells, but not in FLUAV-infected cells.

**Figure 7.**
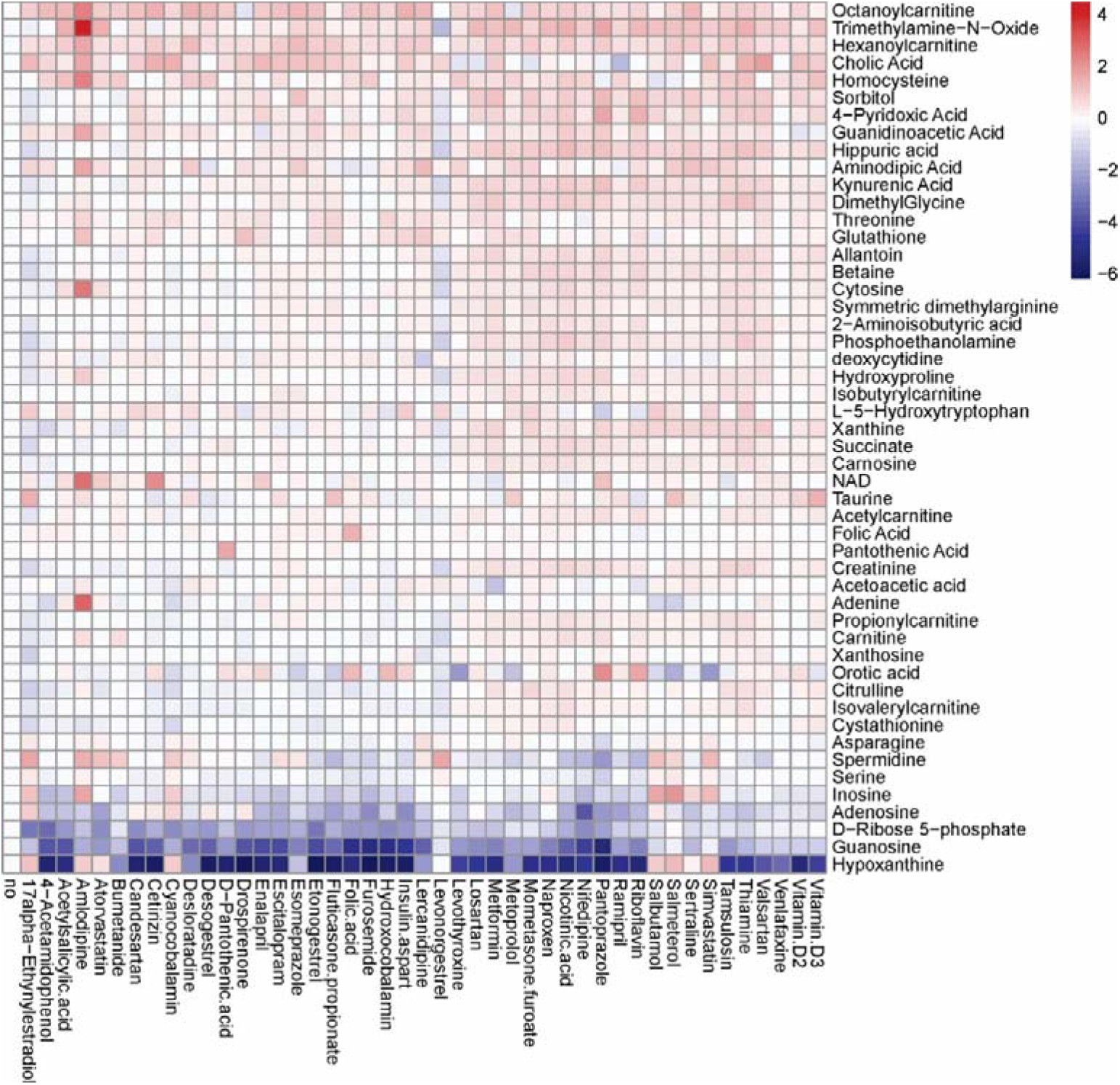
Effect of 45 compounds on metabolism of RPE cells. The cells were treated with 10 µM compounds or remained nontreated. After 24 h, the media were collected and polar metabolites were analysed using LC-MS/MS. A heatmap of 50 most variable metabolites affected by treatment is shown. Rows represent metabolites, columns represent samples. Each cell is coloured according to the log2-transformed and quantile-normalized values of the samples, expressed as fold-change relative to the nontreated control.

**Figure 8.**
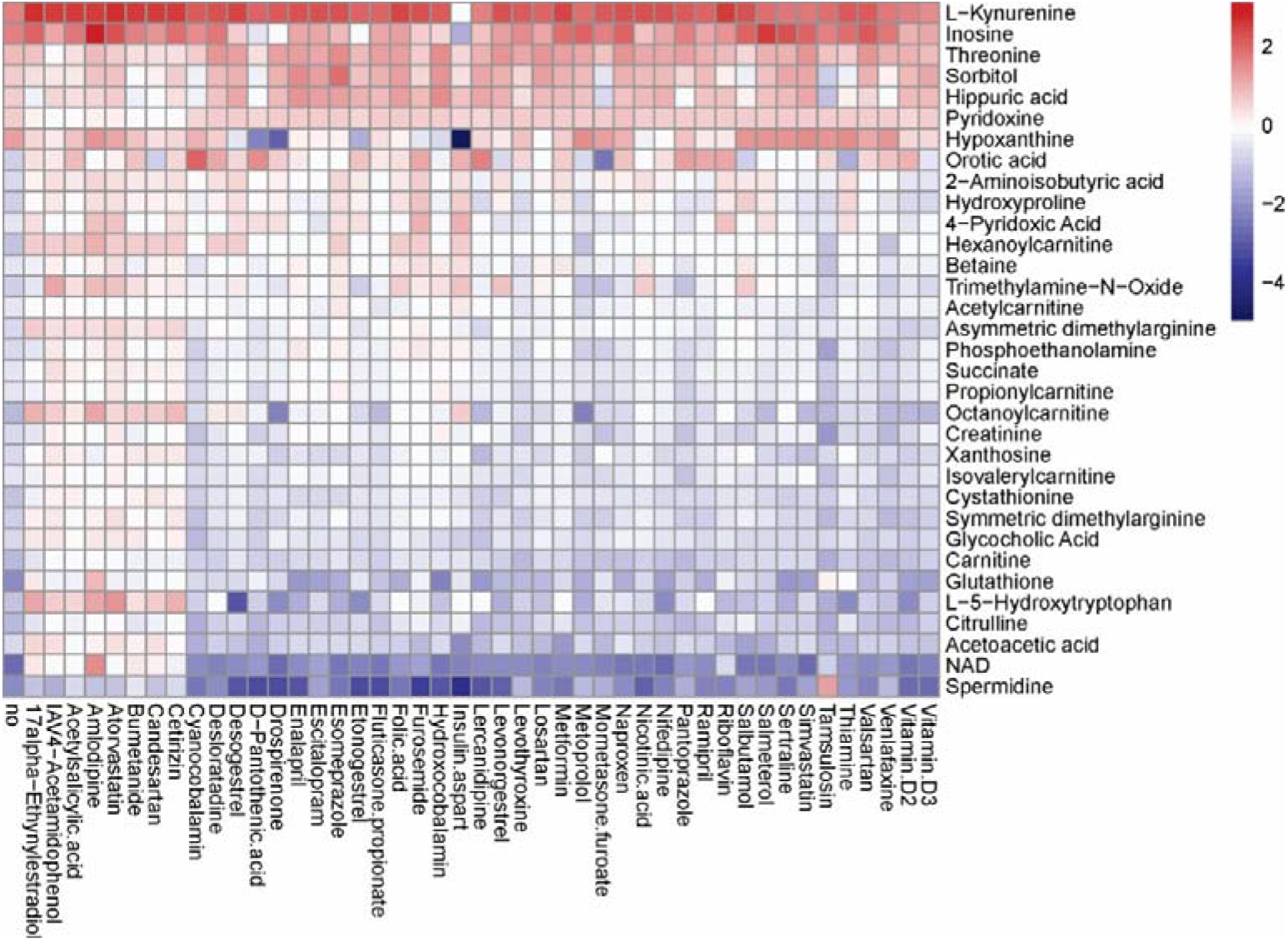
Effect of 45 compounds on metabolism of FLUAV-infected RPE cells. The cells were treated with 10 µM compounds or remained nontreated. After 24 h, the media were collected and polar metabolites were analysed using LC-MS/MS. A heatmap of 50 most variable metabolites affected by treatment is shown. Rows represent metabolites, columns represent samples. Each cell is colored according to the log2-transformed and quantile-normalized values of the samples, expressed as fold-change relative to the nontreated mock-infected control.

Other drugs also deregulated metabolism of different polar molecules in the media of mock or FLUAV-infected cells. 17a-ethynylestradiol, 4-acetamidophenol, acetylsalicylic acid, amlodipine, atorvastatin, bumetanide, candesartan and cetirizin elevated levels of many polar metabolites in the media of infected cells. FLUAV infection itself was found to increase the concentration of L-kynurenine and inosine and lower the concentration of spermidine and NAD, but treatments only moderatly affected the levels of these metabolites. From this, we can see that these commonly prescribed medicines can indeed affect the metabolic acitivity of both mock- and FLUAV-infected cells.

## 4. Discussion

The pharmaceutical industry is one of the most strictly regulated industries in the world, ensuring that medicines approved with marketing authorization are safe and effective, and that the benefits of the drugs outweigh potential risks to the patients. Here, we investigated whether commonly prescribed medicines could affect FLUAV-host cell interactions.

To find the most prescribed medicines, we used the Norwegian Prescription Database, which contains information about medicines dispensed from pharmacies based on prescription from doctors. Altogether, we selected 45 most prescribed medicines. We constructed network of interaction between 45 active compounds of commonly prescribed medicines and host factors, keeping in mind downstream and other physiologic targets. We found several host factors which could affect FLUAV replication within a cell including CXCR4, ALB, HRH1, RHOA, ADRA2B, PNP, and MMAB, which are targeted by niacin, losartan, ramipril, acetylsalicylic acid, thyroxine, valsartan, cetirizine, citalopram, atorvastatin, simvastatin, metoprolol, setraline, candesartan and hydroxocobalam. Furthermore, we were able to uncover compounds that affect the transcription and metabolism of mock- and FLUAV-infected cells including acetylsalicylic acid, atorvastatin, candesartan, and hydroxocobalam. Taken together, we can see that many common prescription drugs are able to modulate FLUAV-host cell interaction.

The current study reports on results using human nonmalignant RPE cells and laboratory H1N1 strain. While this lays the groundwork for a proof-of-concept study, we hope to expand this investigation to cover broader and more applicable models, as well as see how these interactions may change with shifting seasonal flu strains and other viruses. In particular, we are interested in reproducing this work in human primary cells such as macrophages and dendritic cells, which are highly applicable in the context of influenza and other viral infections. In addition, we would like to use other omics techniques, such as genomics, epigenetics, proteomics, phosphoproteomics and lipidomics, to allow identification of signatures associated with adverse reactions of the most prescribed medicines. In addition, we would also like to reproduce this work *in vivo*.

## 5. Conclusions

There are thousands of approved drugs currently being used around the world, and more experimental and investigational drugs are being developed daily. Many of these drugs have side effects, which can only be revealed in clinical studies or long-term post-market monitoring and can be challenging and time-consuming to identify. Here we used an *in silico* and *in vitro* approaches [20] for studying effects of 45 commonly prescribed medicines on FLUAV-host cell interaction. By simultaneously leveraging drug-target interaction studies, drug toxicity/efficacy tests, transcriptomics and metabolomics, we were able to identify markers associated with potential side effects in the context of influenza virus infection. Thus, we have outlined a straightforward, *in silico-in vitro* method to identify hidden cross-over effects of common medications. Overall, we believe that our systems biology approach to identifying possible side effects of medications to common illnesses could be applied broadly in the pharmaceutical industry during development of experimental and investigational drugs.

## Author Contributions

Conceptualization, D.K.; methodology, A.I. and R.Y.; software, A.I.; validation, A.I., R.Y. and E.Z.; formal analysis, A.I.; investigation, A.I., R.Y., H.L., and D.K.; resources, M.B., T.T., V.O., E.Z. and D.K.; data curation, A.I.; writing—original draft preparation, D.K.; writing—review and editing, all authors; visualization, A.I., and D.K.; supervision, M.B., T.T., and D.K.; project administration, D.K.; funding acquisition, D.K. All authors have read and agreed to the published version of the manuscript.

## Institutional Review Board Statement

All experiments with viruses were performed in BSL2 laboratory in compliance with the guidelines of the national authorities under appropriate ethical and safety approvals.

## Informed Consent Statement

Not applicable.

## Data Availability Statement

Data supporting reported results can be requested from the corresponding author.

## Acknowledgments

We thank Hege Ramstad for her help with data analysis.

