## Supplementary material for "Active components of commonly prescribed medicines affect influenza A virus-host cell interaction: a pilot study": Sup tables

**Table S1.** Active compounds of selected drugs, their suppliers and catalogue numbers.

| **Drug** | **CAS** | **MW** | **Formula** | **Cat N** | **Pur., %** | **Supplier** |
| --- | --- | --- | --- | --- | --- | --- |
| 17α-Ethynylestradiol | 57-63-6 | 296 | C20H24O2 | E4876-100MG | ≥98 | Sigma Aldrich |
| 4-Acetamidophenol | 103-90-2 | 151 | C8H9NO2 | 102330050 | 98 | Acros Organics |
| Acetylsalicylic acid | 50-78-2 | 180 | C9H8O4 | AC158180500 | 99 | Acros Organics |
| Amlodipine | 88150-42-9 | 409 | C26H31ClN2O8S | CAYM14838 | ≥98 | Cayman Chemicals |
| Atorvastatin | 134523-03-8 | 559 | C33H35FN2O5 | CAYM10493 | ≥98 | Cayman Chemicals |
| Bumetanide | 28395-03-1 | 364 | C17H20N2O5S | CAYM14630 | ≥98 | Cayman Chemicals |
| Candesartan | 139481-59-7 | 440 | C24H20N6O3 | sc-217825 | ≥98 | Santa Cruz Biotechnology |
| Cetirizin | 83881-52-1 | 389 | C21H27Cl3N2O3 | 89126-50MG | ≥98 | Sigma Aldrich |
| Cyanocobalamin | 68-19-9 | 1355 | C63H88CoN14O14P | DRE-C11798500 |  | LGC Standards |
| Desloratadine | 100643-71-8 | 311 | C19H19ClN2 | CAYM16931 | ≥98 | Cayman Chemicals |
| Desogestrel | 54024-22-5 | 310 | C22H30O | CAYM23651 | ≥95 | Cayman Chemicals |
| D-Pantothenic acid | 79-83-4 | 219 | C9H17NO5 | HY-B0430 | ≥98 | MedChemExpress |
| Drospirenone | 67392-87-104 | 367 | C24H30O3 | CAYM23347 | ≥98 | Cayman Chemicals |
| Enalapril | 75847-73-3 | 376 | C20H28N2O5 | J60750.03 | ≥97 | Alfa Aesar |
| Escitalopram | 128196-01-0 | 324 | C20H21FN2O | CAYM22405 | ≥98 | Cayman Chemicals |
| Esomeprazole | 161973-10-0 | 767 | C34H42MgN6O9S2 | CAYM17326 | ≥95 | Cayman Chemicals |
| Etonogestrel | 54048-10-1 | 324 | C22H28O2 | CAYM21062 | ≥98 | Cayman Chemicals |
| Fluticasone propionate | 80474-14-2 | 445 | C25H31F3O5S | 462101000 | ≥96 | Acros Organics |
| Folic acid | 59-30-3 | 441 | C19H19N7O6 | J62937.06 | ≥97 | Alfa Aesar |
| Furosemide | 54-31-9 | 331 | C12H10ClN2O5S | 448970010 | ≥97 | Acros Organics |
| Hydroxocobalamin | 13422-5 51-0 | 1346 | C62H89CoN13O15P | CAYM24099 | ≥95 | Cayman Chemicals |
| **Drug** | **CAS** | **MW** | **Formula** | **Cat N** | **Purity,  %** | **Supplier** |
| Insulin aspart | 116094-23-6 | 5826 | C256H387N65O79S6 | EPY0000349 |  | LGC Standards |
| Lercanidipine | 132866-11-6 | 612 | C36H41N3O6 | HY-B0612A | 98.5 | MedChemExpress |
| Levonorgestrel | 797-63-7 | 312 | C21H28O2 | CAYM10006 | ≥95 | Cayman Chemicals |
| Levothyroxine | 25416-653 | 817 | C15H12I4NNaO5 | FT48192 | ≥97 | Carbosynth |
| Losartan | 114798-26-4 | 423 | C22H23ClN6O | FL39656 | ≥97 | Carbosynth |
| Metformin | 1115-70-4 | 166 | C4H12ClN5 | sc-202000 | ≥99 | Santa Cruz Biotechnology |
| Metoprolol | 51384-51-1 | 267 | C15H25NO3 | sc-264643 | 97 | Santa Cruz Biotechnology |
| Mometasone furoate | 83919-23-7 | 521 | C27H30Cl2O6 | CAYM21365 | ≥98 | Cayman Chemicals |
| Naproxen | 22204-53-1 | 230 | C14H14O3 | CAYM70290 | ≥99 | Cayman Chemicals |
| Nicotinic acid | 59-67-6 | 123 | C6H5NO2/HOOC5H4N | 128290050 | 99.5 | Acros Organics |
| Nifedipine | 21829-25-4 | 346 | C17H18N2O6 | CAYM11106 | ≥98 | Cayman Chemicals |
| Pantoprazole | 102625-70-7 | 383 | C16H15F2N3O4S | CAYM21345 | ≥98 | Cayman Chemicals |
| Ramipril | 87333-19-5 | 417 | C23H32N2O5 | FC27676 | ≥98 | Cymit Quimica |
| Riboflavin | 83-88-5 | 376 | C17H20N4NaO9P | A11764.14 | 98 | Alfa Aesar |
| Salbutamol | 18559-94-9 | 239 | C13H21NO3 | CAYM21003 | ≥98 | Cayman Chemicals |
| Salmeterol | 89365-50-4 | 416 | C25H37NO4 | HY-14302 | 99.7 | MedChemExpress |
| Sertraline | 79559-97-0 | 306 | C17H18Cl3N | 462190010 | ≥98 | Acros Organics |
| Simvastatin | 79902-63-9 | 419 | C25H38O5 | 458840010 | 98 | Acros Organics |
| Tamsulosin | 106463-17-6 | 445 | C20H29ClN2O5S | CAYM24020 | ≥98 | Cayman Chemicals |
| Thiamine | 67-03-8 | 337 | HC12H17ON4SCl2 | 148990100 | 99 | Acros Organics |
| Valsartan | 137862-53-4 | 436 | C24H29N5O3 | sc-220362 | ≥98 | Santa Cruz Biotechnology |
| Venlafaxine | 99300-78-4 | 277 | C17H27NO2 | HY-B0196A | 98 | MedChemExpress |
| Vitamin D2 | 50-14-6 | 397 | C28H44O | CAYM11791 | ≥98 | Cayman Chemicals |
| Vitamin D3 | 67-97-0 | 385 | C27H44O | CAYM11792 | ≥98 | Cayman Chemicals |

**Table S2.** The most dispensed medicines in Central Norway in 2019 sorted by daily defined dosage (DDD).

| **ATC** | **Active compounds** | **Indications** | **DDD** |
| --- | --- | --- | --- |
| C10AA05 | Atorvastatin | Hypercholesterolemia | 24 480 740 |
| B01AC06 | Acetylsalicylic acid | Pain, fever or inflammation | 16 113 802 |
| C08CA01 | Amlodipine | Hypertension and coronary artery disease | 10 990 638 |
| C09CA06 | Candesartan | Hypertension | 9 404 137 |
| A02BC02 | Pantoprazole | Erosive esophagitis and Zollinger-Ellison syndrome | 8 626 254 |
| R06AE07 | Cetirizine | Hay fever, allergies, angioedema, and urticaria | 8 377 836 |
| C09AA05 | Ramipril | Hypertension and congestive heart failure | 8 034 213 |
| N02BE01 | Paracetamol | Pain and fever | 7 548 361 |
| C10AA01 | Simvastatin | Hypercholesterolemia | 7 117 959 |
| H03AA01 | Levothyroxine sodium | Thyroid hormone deficiency | 6 701 185 |
| G03AA07 | Levonorgestrel | Birth control (in combination with the estrogen ethinylestradiol), emergency birth control | 6 677 552 |
| G03AA07 | Ethinylestradiol | Birth control and treatment of menopausal symptoms in combination with progestins | 6 677 552 |
| B03BB01 | Folic acid | Folate deficiency | 6 598 841 |
| A12AX | Vitamin D2 | Vitamin D deficiency | 5 675 283 |
| A11CC05 | Vitamin D3 (colecalciferol) | Vitamin D deficiency | 5 332 710 |
| R06AX27 | Desloratadine | Allergic rhinitis, nasal congestion | 5 309 712 |
| C07AB02 | Metoprolol | Hypertension and coronary artery disease | 5 264 848 |
| A02BC05 | Esomeprazole | Gastroesophageal reflux disease, erosive esophagitis, duodenal ulcers | 5 152 910 |
| N06AB10 | Escitalopram | Depression, generalized anxiety disorder | 4 728 187 |
| A10BA02 | Metformin | Type 2 diabetes, polycystic ovary syndrome | 4 351 201 |
| G03AC08 | Etonogestrel | Birth control | 4 276 000 |
| M01AE52 | Naproxen | Pain and fever caused by inflammation | 4 031 853 |
| C09CA03 | Valsartan | Hypertension and congestive heart failure | 3 514 084 |
| C09CA01 | Losartan | Hypertension | 3 491 802 |
| G03AC09 | Desogestrel | Birth control and menopausal symptoms | 3 301 519 |
| B03BA03 | Hydroxocobalamin | Vitamin B12 deficiency | 3 182 950 |
| R03AC02 | Salbutamol | Asthma | 3 018 486 |
| A11E | D-Pantothenic acid* | Vitamin B deficiency | 2 872 185 |
| A11E | Thiamine* | Vitamin B deficiency | 2 872 185 |
| A11E | Riboflavin* | Vitamin B deficiency | 2 872 185 |
| A11E | Nicotinic acid* | Vitamin B deficiency | 2 872 185 |
| C03CA01 | Furosemide | Hypertension and edema | 2 864 836 |
| R03AK06 | Fluticasone propionate | Asthma, allergic rhinitis, atopic dermatitis | 2 668 274 |
| C08CA13 | Lercanidipine | Hypertension | 2 667 274 |
| B03BA01 | Cyanocobalamin | Vitamin B12 deficiency | 2 406 614 |
| C03CA02 | Bumetanide | Heart failure | 2 354 487 |
| N06AB06 | Sertraline | Depression | 2 129 315 |
| A10AB05 and A10AD05 | Insulin aspart | Diabetes mellitus type 1 and 2 | 2 101 161 |
| G04CA02 | Tamsulosin | Benign prostatic hyperplasia, kidney stones, acute urinary retention | 2 065 873 |
| R01AD09 | Mometasone furoate | Symptoms in nose caused by allergy or polyps | 1 968 160 |
| N06AX16 | Venlafaxine | Depression, general anxiety disorder | 1 840 761 |
| R03AK06 | Salmeterol | Asthma | 1 833 358 |
| C08CA05 | Nifedipine | Hypertension and angina pectoris | 1 773 040 |
| C09AA02 | Enalapril | Hypertension, diabetic kidney disease and heart failure | 1 770 279 |
| G03AA12 | Drospirenone | Birth control and menopausal symptoms | 1 725 864 |

** B-Tonin, TroBe and nycoplus B-kompleks include d-pantothenic acid (vitamin B5), thiamine (vitamin B1), riboflavin (vitamin B2) or nicotinic acid (vitamin B3).*
